# Isolates from ancient permafrost help to elucidate species boundaries in *Acanthamoeba castellanii* complex (Amoebozoa: Discosea)

**DOI:** 10.1101/755348

**Authors:** Stas Malavin, Lyubov Shmakova

## Abstract

*Acanthamoeba castellanii* species complex (genotype T4) comprises of more than ten species with unclear synonymy. Its molecular phylogeny has several conflicts with published morphological data. In this paper, we analyze morphometric traits and temperature preferences in six new strains belonging to *A. castellanii* complex isolated from Arctic permafrost in the framework of molecular phylogeny. This integrative approach allows us to cross-link genotypic and phenotypic variability and identify species-level boundaries inside the complex. We also analyze previously known and newly found discrepancies between the nuclear and mitochondrial gene-based phylogenies. We hypothesize that one reason for these discrepancies may be the intragenomic polymorphism of ribosomal RNA genes.

## Introduction

*Acanthamoeba* is a protist genus belonging to the supergroup Amoebozoa. It has very characteristic angular, or stellate, cysts with an envelope consisting of two distinct layers (Fig. 1K, L, M) and fine tapered hyaline projections in the active forms – the so-called acanthopodia (Fig. 1A, B, D, E). It is common in soils and aquatic environments all around the world (Geisen et al., 2014). The genus is also known as an opportunistic pathogen causing amoebic keratitis and encephalitis in humans (Khan, 2005; Maycock and Jayaswal, 2016). Due to a well-established method of axenic cultivation, it has become a popular model in biochemical and cell structure studies (Siddiqui and Khan, 2012). *Acanthamoeba* was among the first organisms whose nuclear small subunit ribosomal RNA gene (18S rDNA) sequence was determined (Gunderson and Sogin, 1986). Its genome is assembled and annotated (Clarke et al., 2013), and several other draft assemblies are present in GenBank.

**Figure 1.**
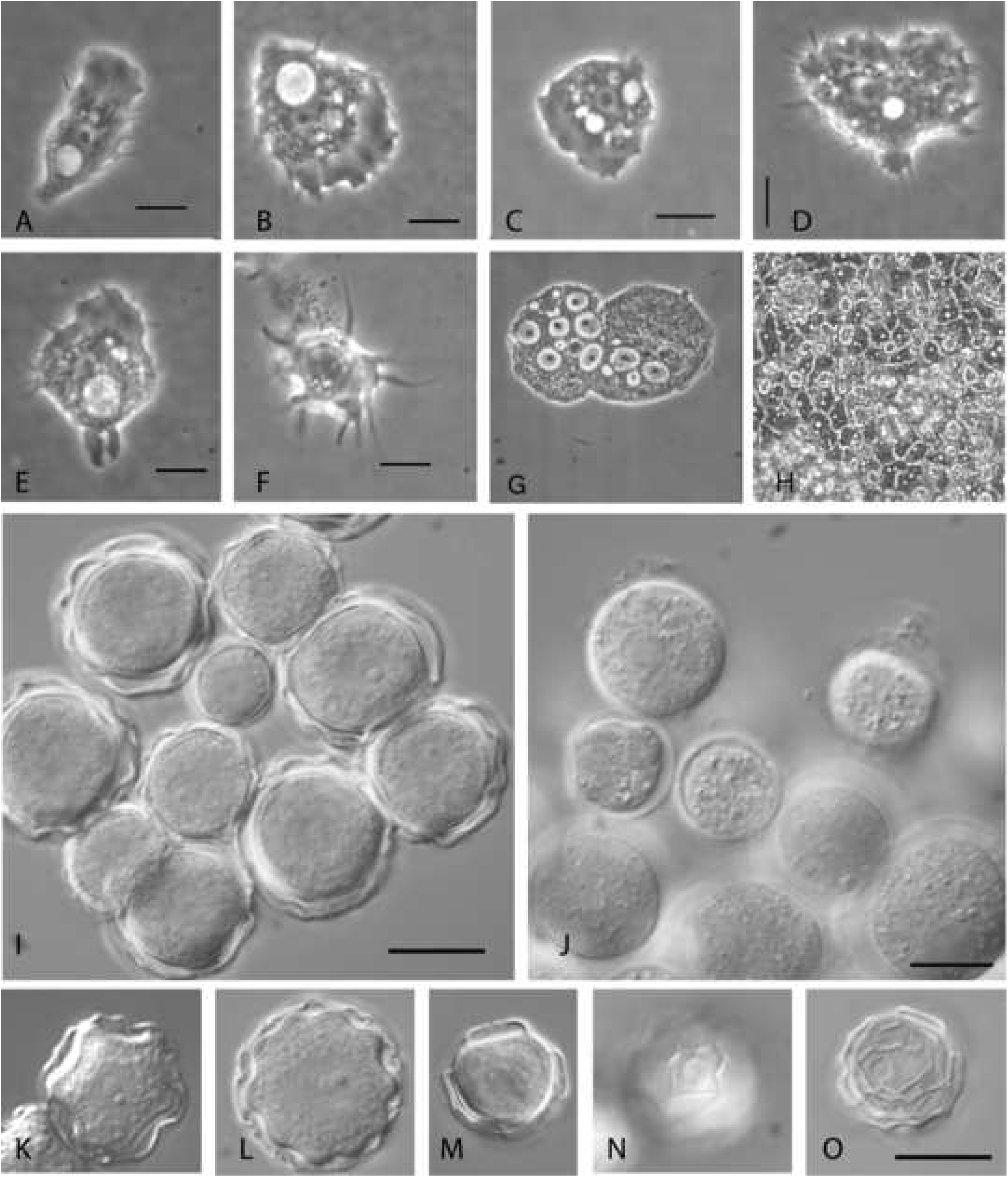
**A–E** Locomotor forms: A – SCL-14-3; B – SCL-14-3; C – SCL-14-12, D – SCL-am9; E – SCL-14-3; **F** Floating form, SCL-am8; **G** – multinucleate trophozoite, SCL- am9; **H** Monolayer, SCL-14-12; **I, K–N** Cysts, various strains; **J** Pseudocysts, SCL- am8; **O** Empty cyst, SCL-am8.

Despite such a rich history of study (or partially because of it), the system of the genus is surprisingly tangled. The list of described species includes around 30 names. For several of them, the synonymy is apparent from allozyme and molecular data and could also be suspected for some others. Since the beginning of rDNA phylogeny era, several ribotypes (traditionally called “genotypes” in *Acanthamoeba* literature) have been described based on a threshold in genetic distances and characteristic signatures in genus-specific hypervariable regions of the 18S gene. These types were designated T1–T22 (Corsaro and Venditti, 2018; Tice et al., 2016). While some of the types directly correspond to formally described species, others include type strains of several species. This particularly concerns type T4, the most frequently isolated one in both clinics and nature (Maciver et al., 2013). Internally, type T4 is also heterogeneous, which has been noted since the beginning of comparative studies of the 18S rDNA in *Acanthamoeba* (Fuerst et al., 2015; Gast et al., 1996), though genetic distances between clusters inside T4 are smaller than those between this and the other types (Stothard et al., 1998). Here belong the type strains of most of the species described on the basis of morphology, including the generic type, *Acanthamoeba castellanii* (Douglas, 1930) and thoroughly redescribed by Page (1967) *Acanthamoeba polyphaga* (Puschkarew, 1913). Due to the plastic nature of characters by which T4 species are differentiated and unclear synonymy, some authors prefer to designate this type “*A. castellanii* species complex” (Corsaro et al., 2017; Kong, 2009). Therefore, reconciling molecular with morphological data inside T4 is essential for the system of the genus as a whole, which means establishing a genus-specific species concept integrating phenotypic and genotypic taxonomic markers (Boenigk et al., 2012).

We isolated six new *Acanthamoeba* strains from Arctic permafrost sediments, a specific environment, where they have been buried and trapped in a dormant but viable state. It is shown that in cases when freezing occurs in parallel to sedimentation (which can be proven by geological data), the age of embedded cells corresponds to the age of the permafrost sediments, as both transportation and vital activity is impossible at temperatures below 10°C (Rivkina *et al*., 2000; Gilichinsky & Rivkina, 2011). Our strains were isolated from the sediments dated from the late Pleistocene to early Holocene. Previously, other eukaryotes, including ciliates (Shatilovich et al., 2015), amoebozoans (Shmakova et al., 2016, 2018), flagellates (Shatilovich et al., 2009; Stoupin et al., 2012), green algae (Vishnivetskaya, 2009), fungi (Kochkina et al., 2012), and nematodes (Shatilovich et al., 2018) were isolated from the sediments of similar age. Even a higher plant belonging to the genus *Silene* was regenerated from the tissue of a seed found in the excavated borrowing of a ground squirrel, although the attempts to grow those seeds failed (Yashina et al., 2012).

All isolated *Acanthamoeba* strains belong to ribotype T4. To precisely identify the taxonomic position of the strains, we additionally sequenced nearly full-length 16S (mitochondrial) gene and the barcode region of cytochrome oxidase subunit I (Cox1) gene. Further, we performed morphometrics of all life cycle stages, including cysts formed in different conditions. We also studied the growth of isolated strains at different temperatures. By integrating these data, we identified a species-level boundary within the studied range of variation. Putting morphometrics and ecophysiology of several strains into a broader (almost genus-wide) phylogenetic framework allowed us to figure out which level of genetic heterogeneity corresponds to the species-level difference in phenotypes. Additionally, we analyzed previously known and newly identified discrepancies between the phylogenies based on different genes and hypothesized their origin.

## Material and Methods

### Sampling and strain isolation

Samples for subsequent enrichment cultivation of amoebae were taken from the intact inner part of the permafrost cores. The cores were obtained in the Gydan Peninsula (Western Siberia), Kolyma Lowland, and Bykovsky Peninsula (Eastern Siberia) by a special drilling technique allowing for the sterility of the sample, as described elsewhere (Gilichinskiy et al., 1989; Shi et al., 1997). For the map of the borehole locations see Supplementary Fig. 1. The cores were kept frozen during all steps of transportation and laboratory processing, which completely prevents the contamination of the inner material by modern microorganisms (Juck *et al*., 2005; Gilichinsky & Rivkina, 2011).

Samples of about 1 cm^3^ were taken from the cores in a laminar flow hood using a sterile scalpel and put into sterile Petri dishes filled with autoclaved mineral Prescott and James medium (Page, 1988). No bacterial culture was added, as reproducing viable bacteria from permafrost soil formed sufficient food supply for the emerging amoebae. The plates have been checked daily under a phase-contrast inverted microscope, and observed amoebas transferred individually by a tapering Pasteur pipette into new plates, from which clonal cultures were subsequently established.

### Culturing and encystment

Strains were maintained as monoclonal bacterized cultures (the bacterial community originated from initial permafrost sample) on 1.5% non-nutrient agar plates with 0,01% cerophyl infusion overlay (Page, 1988) at 18°C. Cultures were passaged monthly. Cysts formed in such cultures without any special induction one to two weeks after plating.

For axenization, cyst-rich cultures were washed from the agar layer, collected by centrifugation at 1,000×g for 1 min, and treated with 1M HCl for 24 hours. The remaining cysts were then washed thrice with autoclaved distilled water and incubated in sterile proteose-yeast-glucose (PYG) medium (Neff et al., 1964) with 100 U mL^-1^ of penicillin and 0.1 mg mL^-1^ of streptomycin (Sigma-Aldrich). Axenic cultures were maintained in 25 cm^2^ culture flasks in the same medium at 18°C and passaged every two weeks.

Three conditions of encystment were tested: “bacterized culture,” “sparse axenic culture,” and “dense axenic culture.” A typical bacterized culture represented the first. Axenic cysts were obtained from cultures grown in 3 mL PYG on 60 mm plates to the monolayer stage. The “sparse axenic culture” conditions were achieved as follows: a subculture of 200 μL was transferred into 3 mL Prescott and James (PJ) mineral medium (Page, 1988) and left for several days. The rest of the cells were scraped and centrifuged at 1,000×g for 1 min, washed with PJ, and left in 3 mL PJ for several days representing the “dense axenic culture” conditions. From now on, we refer to the cysts formed in “bacterized culture” conditions as “bacterized cysts”; to those formed in “sparse axenic culture” conditions as “sparse axenic cysts”; to the cysts formed in “dense axenic culture” conditions as “dense axenic cysts.” Pseudocysts were obtained from axenic cultures. Cells grown at 18°C in 60 mm Petri dishes (3 mL PYG) up to the monolayer stage were washed once with PJ, collected by centrifugation, and put into 3 mL 5% dimethyl sulfoxide (DMSO) solution in PJ (Kliescikova et al., 2011). To test the viability of the obtained pseudocysts, they were put into fresh PYG medium, either immediately or after being completely dry during a week.

### Microscopy and measurements

Living trophozoites and cysts were observed and filmed either on slides with petroleum jelly sealing using Leica DM2500 microscope with phase and differential contrast optics or directly in culture using Zeiss Axiostar Plus microscope with phase contrast optics and a water-immersion objective. Nikon DS-Fi3 and Watec WAT-221S digital cameras were used for filming in the first and the second case, correspondingly.

Measurements were performed on a monitor with a calibrated ruler using pictures obtained with a Zeiss Axiostar Plus microscope equipped with a Watec WAT-221S camera. In the case of trophozoites, locomotor forms and rounded cells were measured. The measured parameters in locomotor forms included (1) the longest dimension (“the length”), (2) the dimension perpendicular to the longest and going through the nucleus (“the width”), (3) the diameter of the nucleus, (4) the diameter of the nucleolus, (5) the maximum width of the hyaloplasm crescent at the anterior edge of the cell. For the rounded cells, cysts, and pseudocysts, the diameter was registered. For the cysts, we also counted the number of ostioles. The counting was done without any special staining using phase contrast optics caused the ostiole complex to appear glowing, which in the case of our strains was found to allow practically unambiguous identification of the ostiole number. The cyst measurements were performed across all three conditions tested (see Culturing and encystment).

### Statistical analysis of morphometric characters

All calculations were performed in R v. 3.5.2 (R Core Team, 2019). We assumed no normality assumptions for all variables. For a measure of central tendency, we calculated medians and expressed variation as a median absolute deviation (as implemented in package *stats*; scale factor 1.4826) or a range. Values exceeding three interquartile range around the median were removed as outliers. The statistical significance of median differences was assessed with Kruskal-Wallis rank-sum test (package *stats*) and non-parametric (“permutational”) analysis of variance (PERMANOVA), as implemented in the function *adonis*, package *vegan* v. 2.5-3 (Oksanen et al., 2019). Between-groups pairwise Wilcoxon rank-sum tests with Benjamini & Yekutieli correction for multiple comparisons (package *stats*) was used for the post-hoc comparison. Clustering of the morphometric data was done using the neighbor-joining algorithm on euclidean distances, as implemented in *dist* and *hclust* functions.

### Sequence acquisition

DNA was extracted from axenic cultures grown in 60-mm Petri dishes to the stationary phase. Cells were scraped from the plastic and collected by centrifugation as described above. The pellets were washed thrice with autoclaved distilled water and finally resuspended in 100 μL of it. This suspension was subjected to the DNA isolation by ExtractDNA Blood kit (Proteinase K plus SDS based lysis and silica-membrane purification; Evrogen, Russia), according to the manufacturer protocol.

The nuclear SSU rRNA gene (18S), mitochondrial SSU rRNA gene (16S), and the mitochondrially encoded cytochrome c oxidase subunit I gene (Cox1) were amplified, purified by agarose gel electrophoresis, and sequenced on an automated Sanger sequencer in Evrogen Co. (Russia). Details of PCR and sequencing may be found in the Supplementary Material.

GenBank accession numbers for the newly obtained sequences are as follows. 16S: MK100243–MK100248. Cox1: MK105577–MK105582. 18S: MK124583–MK124588.

### Phylogenetic analysis

For phylogenetic inference, all available good-quality (nearly) full-length sequences were downloaded from GenBank. MAFFT v. 7.307 L-INS-i algorithm (Katoh and Standley, 2013) was used to align sequences. For 18S, hypervariable regions of the alignment were allocated into a separate partition, where gaps were coded as the fifth state (Simmons and Ochoterena, 2000). A phylogeny was reconstructed by Bayesian analysis of posterior probabilities (Bayesian inference, BI) in MrBayes v.3.2.6 (Ronquist et al., 2012) and maximum likelihood (ML) approach in IQTree v.1.6.6 (Nguyen et al., 2015). The detailed description of the phylogenetic analyses may be found in Supplementary Material.

### Growth at different temperatures

We tested the growth of the isolated strains at 4°C, 11°C, 18°C, 25°C, 32°C, 37°C, 40°C, and 42°C (Δ = 1°C for all). At 4–32°C, only axenic cultures were tested, while to 37–42°C, both axenic and bacterized cultures were exposed. Preliminarily, all temperatures were checked with 40-mm plates (1 mL PYG) in several replicates, to test if growth is possible at all. Axenic cultures for the tests were obtained by transferring 50 μL of a 5-day culture. In the case of bacterized cultures, test plates were inoculated with 50 μL of a two-week culture. This assay was carried out for one week; the cultures were checked daily.

For the temperatures, at which the strains have demonstrated growth or at least persisted successfully, the growth rate was estimated quantitatively. 12-well culture plates, 2 mL of PYG per well, were inoculated with a growing axenic culture as described above. Each strain was tested in two to three replicates per temperature level. The assays were conducted until detached rounded cells started to prevail in the culture. Cells were counted in a hemocytometer each 24 h (each 12 h at 32°C). At 4°C, the experiment lasted for a month, and samples for counting were taken weekly, as observations showed no discernible increase of the cell number. The culture growth was approximated by a logistic curve 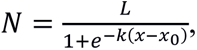 where *L* is the asymptote and *k* – the logistic growth rate. The approximation was done by Levenberg-Marquardt nonlinear least-squares algorithm as implemented in MINIPACK library interfaced through *minpack.lm* package v.1.2-1 (Elzhov et al., 2016) for R v.3.5.2.

## Results

### Isolated strains

In total, we have obtained ten *Acanthamoeba* isolates from around 400 studied permafrost sediment samples from East and West Siberia. We have succeeded in cloning and axenization of six out of ten isolates. The location of the sampling sites is shown in Supplementary Fig. 1. For the essential characteristics of the samples, see Tab. 1.

### Microscopy observations

All obtained strains feature typical morphological characters of the genus, i.e., acanthopodia in trophozoites and angular shape of the endocyst, and belong to the morphotype II following Pussard and Pons (1977).

#### Trophozoites

Trophozoites of all strains readily develop locomotor forms when transferred to fresh medium or the slide. The locomotor form is flattened, with a distinct frontal hyaline area usually occupying about one-fourth of the cell length (Fig. 1A–E). Typically, it is elongated in the direction of movement, with a width-to-length ratio of about 0.6 (Fig. 1A, B). Occasionally, we have observed locomotor forms with the width equal to or even greater than length (Fig. 1C, D). Such forms have been encountered more often in strains SCL-14-2, SCL- 14-3, SCL-14-12, and are generally more typical for larger cells. Acanthopodia are absent on the leading lobopodium in locomotor forms, though the hyaline crescent edge has rarely been even for a long time, regularly demonstrating undulations and protrusions. In cells moving irregularly, acanthopodia are present on lobopodia also. At the beginning of an active movement or while changing its way, trophozoites demonstrate sudden extensions of the lobopodium in a new direction similar to the eruptive movements of heterolobose amoebae.

Generally, one vesicular nucleus with a single nucleolus is situated in the central part of the cell in locomotor forms. At temperatures 25 to 32 °C, we have observed multinucleate trophozoites (Fig. 1G). Those have always been bigger than the median, while nucleoli have been smaller than in normal cells (Fig. 1G).

A single contractile vacuole is usually situated in the rear part of the moving cell (Fig. 1A, B). In axenic cultures, the cytoplasm is strongly vacuolated, while in bacterized cultures, the number of vacuoles is found to be considerably lower. A bulbous uroid often appeared in locomotor forms (Fig. 1A, C, D), sometimes with adhesive filaments (Fig. 1D). Sporadically, we observed a sizeable goblet-like food cup at the rear end of the moving cell (Fig. 1E), identically to what was described and figured by Page (1967). This formation was considerably more often found in strain SCL-14-12.

Floating forms have been rarely observed in bacterized cultures. The floating form represents a central body mass with 15–20 radiating pseudopodia up to 15 μm long (Fig. 1F). In axenic cultures grown in plates or culture flasks without agitation, amoebae form a characteristic monolayer on the surface of the substrate (Fig. 1H). The “closeness” of the monolayer depends on the temperature and adhesive properties of the plastic and differs across the vessel area (Fig. 1H demonstrates the “ideal” case). After the monolayer reaches its closeness limit, cells start to appear above it massively. This scenario is followed by all studied strains but SCL-14-12, in which cells begin to form compact clumps above the surface from the early stages of the culture growth, in parallel to the monolayer formation.

When the culture is roughly shaken or centrifuged, trophozoites appear as rounded cells without lobopodia and acanthopodia. Such cells demonstrate the internal movement of organelles, though no cell movements can be observed for some time. Similar rounded cells also start to appear soon after an axenic culture reaches its growth limit. In this case, most of them probably die in some time, as the final number of cysts in such cultures is much lower than the number of rounded cells observed at the end of growth.

#### Cysts

All isolated strains readily encyst in laboratory cultures. In bacterized cultures, all or at least most of the amoebae form cysts in about two weeks after transfer. In axenic cultures, we obtained cysts by transferring amoebae into either the Neff’s encystment medium (Neff et al., 1964) or merely starving PJ medium. When starting to encyst, cells form plaques.

Mature cysts of all strains have well-defined endo- and ectocyst, the former folded and rippled, the latter smoothly polyhedral. Beneath the endocyst, the characteristic layer of oriented granules is always apparent (Fig. 1I, K, L, M). The number of endocyst corners usually vary from 3 to 7 (Fig. 1I, K, M, O), but cysts with 8 and more corners have also been encountered (Fig. 1L). The majority of ostioles lay in one plane. Opperculi do not protrude above the ectocyst level. Dense axenic cysts usually have less prominent endocyst corners than sparse axenic cysts and bacterized culture cysts. In the latter two, a considerable share of cysts has smooth endocyst without any defined corners (Fig. 1I).

#### Pseudocysts

The pseudocyst formation is total only at DMSO concentrations not lower than 5% and when the DMSO-to-cells ratio is not less than the PYG-to-cells ratio during culture growth. In such conditions, all cells transform into pseudocysts in 12 hours. Both dried and non-dried pseudocysts demonstrate almost total viability being put into PYG; in both cases, trophozoites appear in no more than a day.

Pseudocysts are rounded cells with a fine smooth envelope well seen under both phase- and DIC-contrast (Fig. 1J). The nucleus is situated in the central area of the cell; the nucleolus is slightly smaller than in trophozoites. The cytoplasm is evenly filled with refractile granules. The size of the granules in pseudocysts is bigger than that in cysts. No granular layer beneath the envelope characteristic for cysts may be observed.

### Molecular phylogeny

#### 18S

The inspection of the preliminary automatic alignment of the T4 18S sequences has revealed six hypervariable regions corresponding to the parts of the third, fifth, sixth, ninth, and eleventh hypervariable (“expansion”) regions identified by Gast et al. (1996). Two of those six regions are parts of the diagnostic fragment 3 (DF3) (Booton et al., 2002). The DF3 region is used to distinguish sequence types (Booton et al., 2009, 2005, 2002), but we have found it sufficient to discriminate between T4 subtypes also, except for some rare cases of closely related sequences. The alignment of concatenated hypervariable regions has yielded easily discernible patterns (Supplementary Fig. 3) and has, first of all, demonstrated many several-nucleotide-long indels present in many sequences which by any chance could not be attributed to the alignment method.

Both BI and ML analyses produce seven subtypes, T4_A_ through T4_E_ and T4_Neff_, previously mentioned by Fuerst (2014) and Fuerst et al. (2015), further referred to as A–E and Neff. All but A and B receive maximum support in both analyses; B is highly supported (0.9) and A moderately supported (0.78) in the BI analysis. At the same time, the ML and BI trees display one difference in their topologies (Fig. 2). While in both trees the major division lay between the clades A+B and C+F+E, in the ML tree, subtype D appear sister to the A+B clade and subtype Neff sister to subtype E. In the BI tree, vice versa, subtype D is sister to subtype E and subtype Neff sister to the A+B clade.

**Figure 2.**
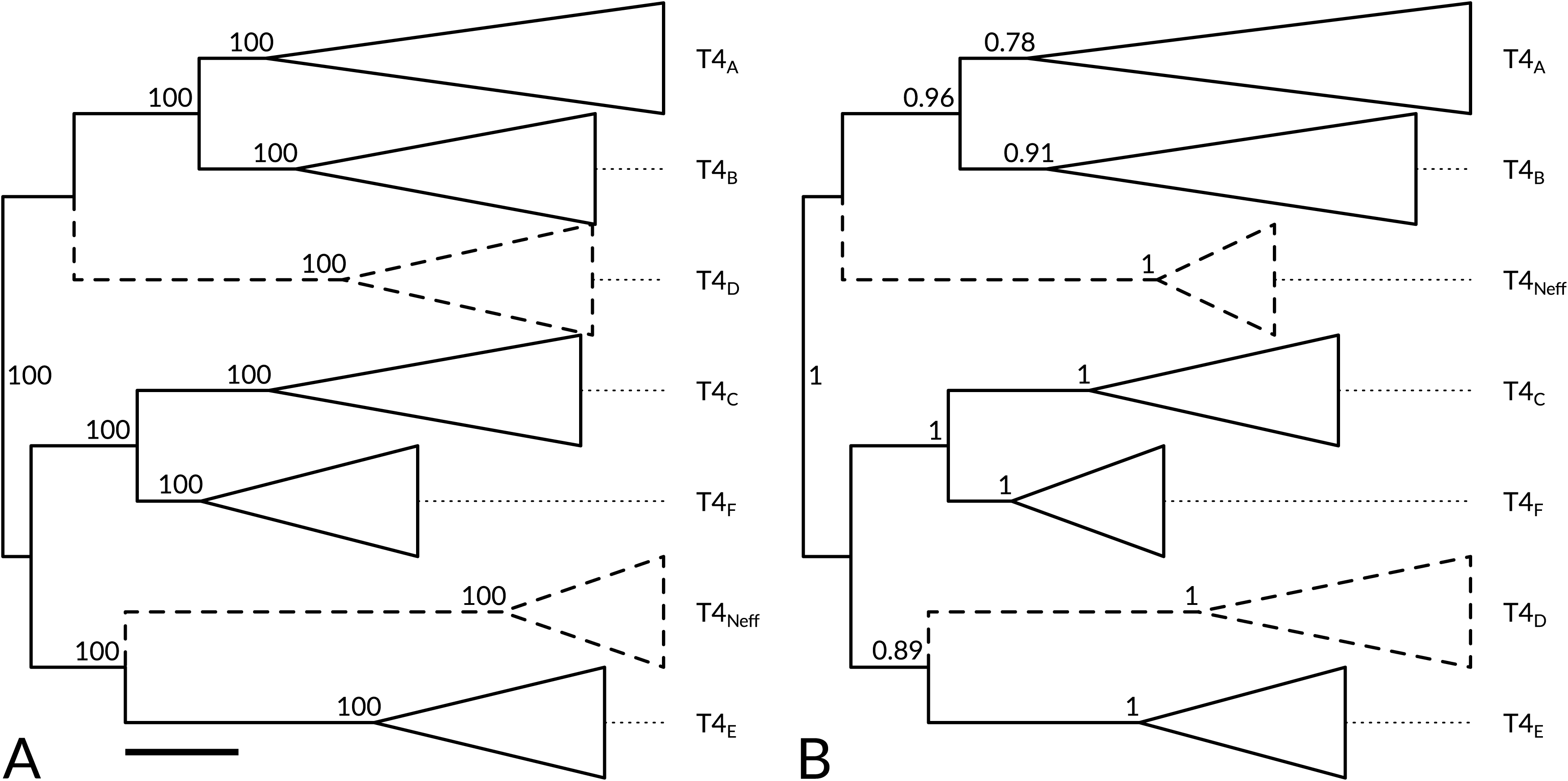
ML (**A**) and BI (**B**) 18S SSU trees of *Acanthamoeba* T4 genotype showing seven subtypes. Branch supports are given as bootstrap percentage / Bayesian posterior probability. The trees are rooted at midpoint. Clades with variable position are dashed. **Scale bar**: 0.01 substitution per site for both A and B.

Several subclusters have got high support in both ML and BI analyses inside clades A and B (Fig. 2). From now on, we refer to them using well-known and long-studied strains they include, for which 16S sequences are also available. This is done for the sake of descriptiveness and does not imply that these strains are somehow “characteristic” of the clusters being described. In clade A, those are “Castellani,” “Galka,” “Haas,” “Rawdon,” “Garcia” clades, and one other, not discussed further. In clade B, those are “Ma,” “Diamond,” “V017” clades, and one other.

In T4_A_, the topology of the Bayesian tree differs from that of the ML tree (Fig. 3) chiefly by the position of “Haas” clade, which is sister to the cluster containing “Galka” and “Castellani” clades instead of being the deepest T4_A_ branch as in the ML tree, and the position of “Garcia” clade placed inside “Rawdon” clade instead of being the second deepest branch as in the ML tree.

**Figure 3.**
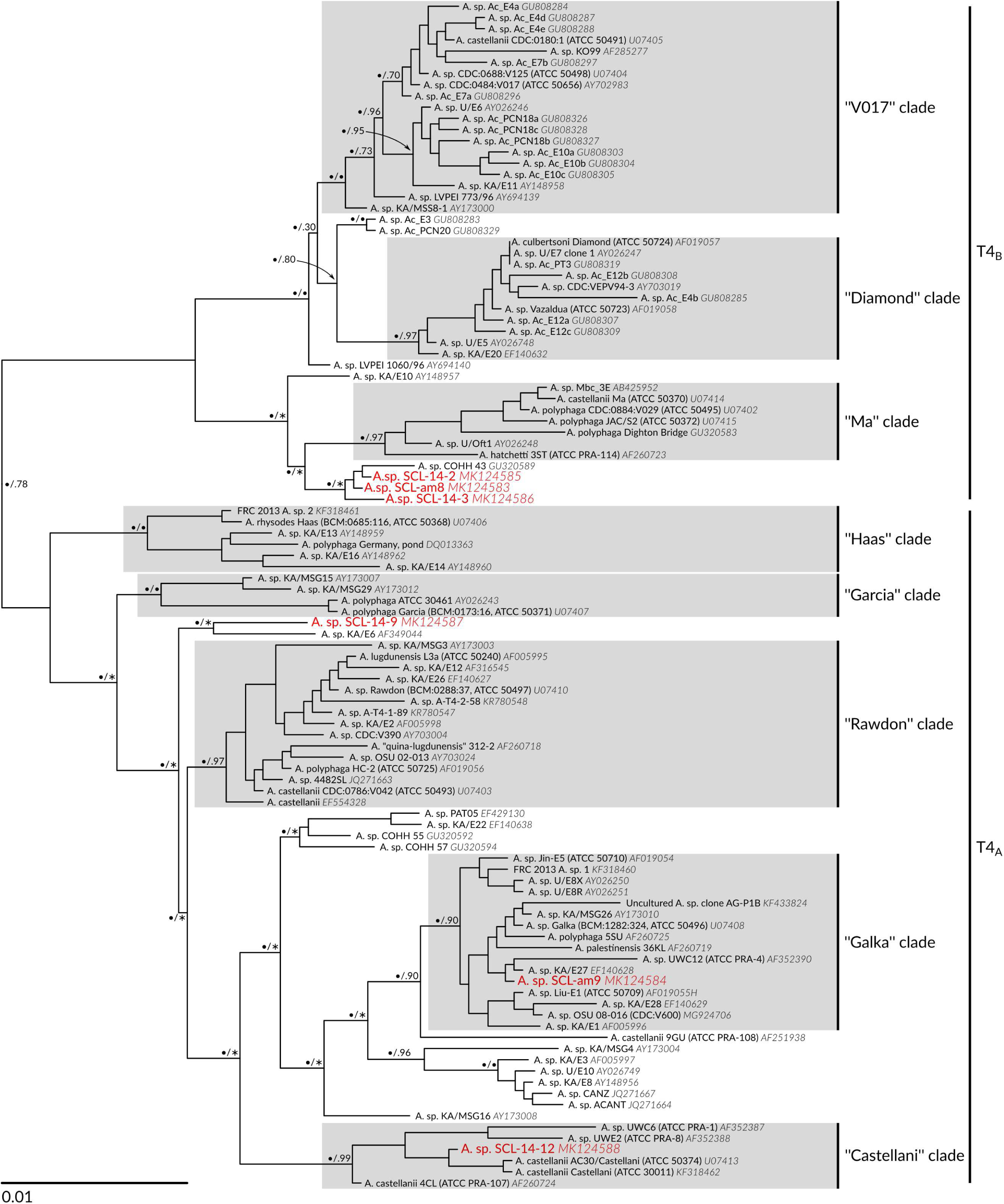
18S SSU ML tree of T4A and T4B clades (a subtree of Fig. 5A, expanded). Bayesian tree gave similar topology (see text). Branch supports are given as bootstrap percentage / posterior probability. • – 100% or 1.00 support; ∗ – clade not observed. Support values inside the clades are omitted. The tree is rooted at midpoint.

Strains SCL-am8, SCL-14-2, and SCL-14-3 branch together in the ML tree (Fig. 3) and group with strain COHH 43 from marine sediments of Mt Hope Bay, MA, USA (Gast et al., 2011) as a distinct clade sister to “Ma” clade of T4_B_. In the BI tree, the permafrost strains and COHH 43 branch before “Ma” clade. Other differences between the BI and ML tree topologies in T4_B_ involved terminal branching only. Three other permafrost strains grouped within T4_A_ clade. Strain SCL-14-9 is relatively distant from the other strains except for one clinical isolate KA/E6 from Korea (Liu et al., 2006); their position in the tree is unstable. Strain SCL-am9 is firmly placed into “Galka” clade, and strain SCL-14-12 – into “Castellani” clade.

#### 16S

The analysis of the 16S alignment confidently recovers all the previously established 18S types for which 16S sequences are available (Fig. 4). We thus refer to them further without mentioning in the framework of which analysis are they inferred. Type T4 is moderately supported (94%) in ML-analysis but has posterior probability of 1 in BI. T2/6 type is sister to T4 type; the next-closest is T5 followed by T3/T11 clade, though this topology as a whole appears moderately supported. Types T7–T9 (morphogroup I by Pussard and Pons (1977) form a very distinct clade with full support.

**Figure 4.**
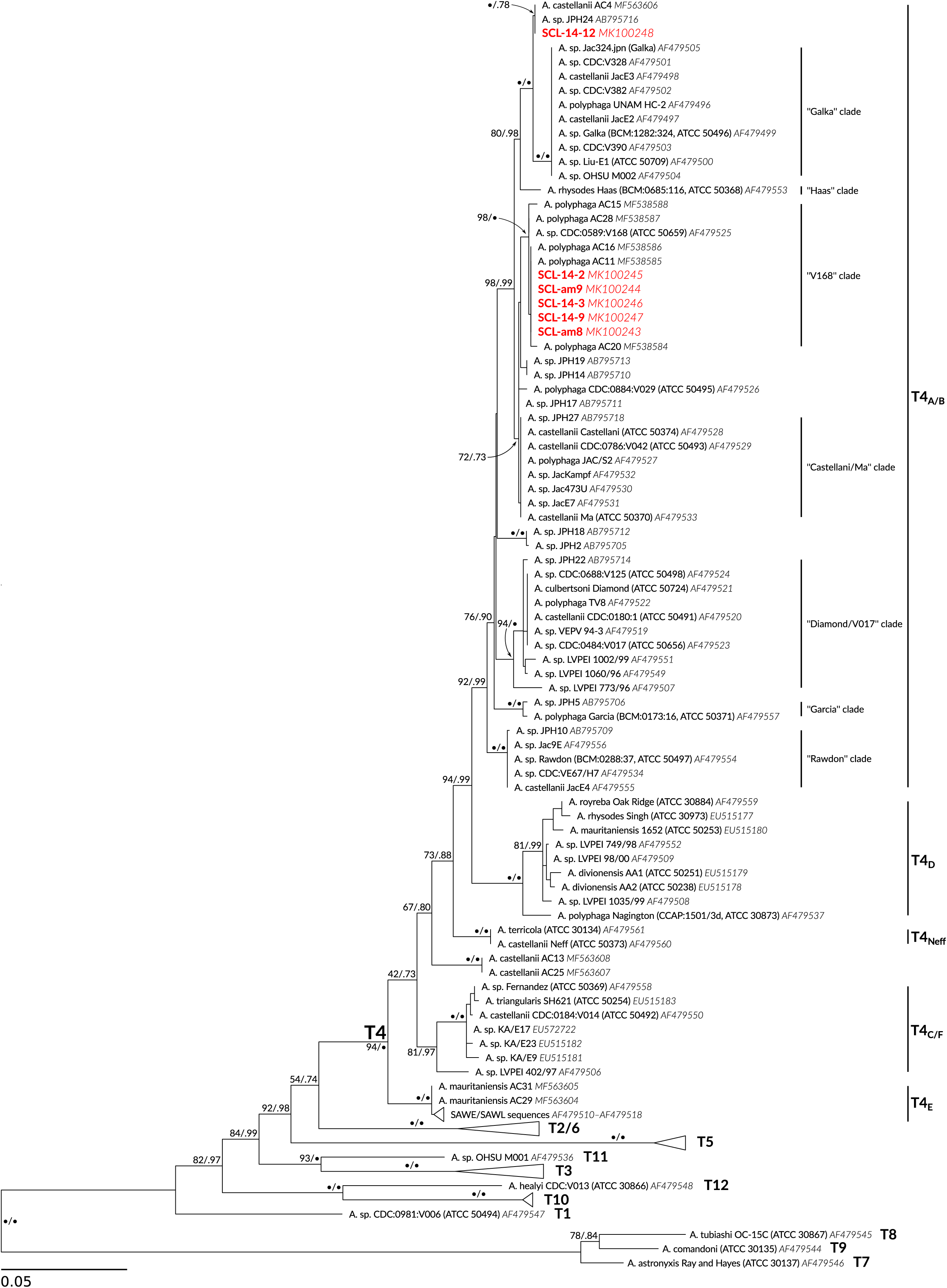
16S SSU ML tree of *Acanthamoeba* T4 genotype. BA tree gave the same topology. Branch supports are given as ML bootstrap percentage / Bayesian posterior probability. • – 100% or 1.00 support. Support values near tree tips are omitted. The tree is rooted at midpoint.

The internal structure of 16S-T4 clade partially reflects that of 18S-T4. D, E, and Neff subtypes appear monophyletic with full support. At the same time, subtypes A and B, as well as C and F, are not separate. Subtype E is the most distant from A/B clade, followed by C/F clade, then subtype Neff. Subtype D is the closest to A/B clade. The 16S tree thus corroborates the 18S one inferred with the ML approach, besides the A/B and C/F fusion.

The blended 16S A/B clade, like its 18S separate counterparts, is internally structured (Fig. 4). Galka, Rawdon, and Garcia strains from 18S-T4_A_ also hit highly supported clusters. Strain Haas does not fall into any cluster but is itself steadily separated. At the same time, strains Castellani and Ma have very similar 16S sequences and form a supported cluster, together with some other strains. The same concerns strains Diamond and CDC:V017.

Surprisingly, all strains from permafrost except SCL-14-12 have identical 16S sequences, also identical to those of strains AC11 and AC16 from India (Megha et al., 2018). In addition to AC15, AC20, AC28, and CDC:V168 (18S-T4_A_), they form a highly-supported cluster inside T4. SCL-14-12 groups together with Indian AC4 (Megha et al., 2018) and JPH24 from Japan (Rahman et al., 2013) (18S-T4_B_) as a sister clade to a group containing strains Galka, HC-2, CDC:V390, Liu-E1 (all 18S-T4_A_), with absolute support. Indian strains AC13 and AC25 reveal rather divergent sequences, probably corresponding to a not yet recovered T4 subtype. Unfortunately, their 18S sequences in GenBank are not full, so it is not possible to either confirm or deny this hypothesis by now.

#### Cox1

As with the 16S gene, strains belonging to Rns D, E, and Neff subtypes fall into different highly supported clusters, while those within Rns A and B, as well as C and F, form fused clusters (Fig. 5). Clade D groups together with clades C/F, E, and Neff. This subdivision is not supported by ML analysis, though gained 0.97 posterior probability. Inside the C/F/Neff half of the tree, no subtype relationships are confidently resolved.

**Figure 5.**
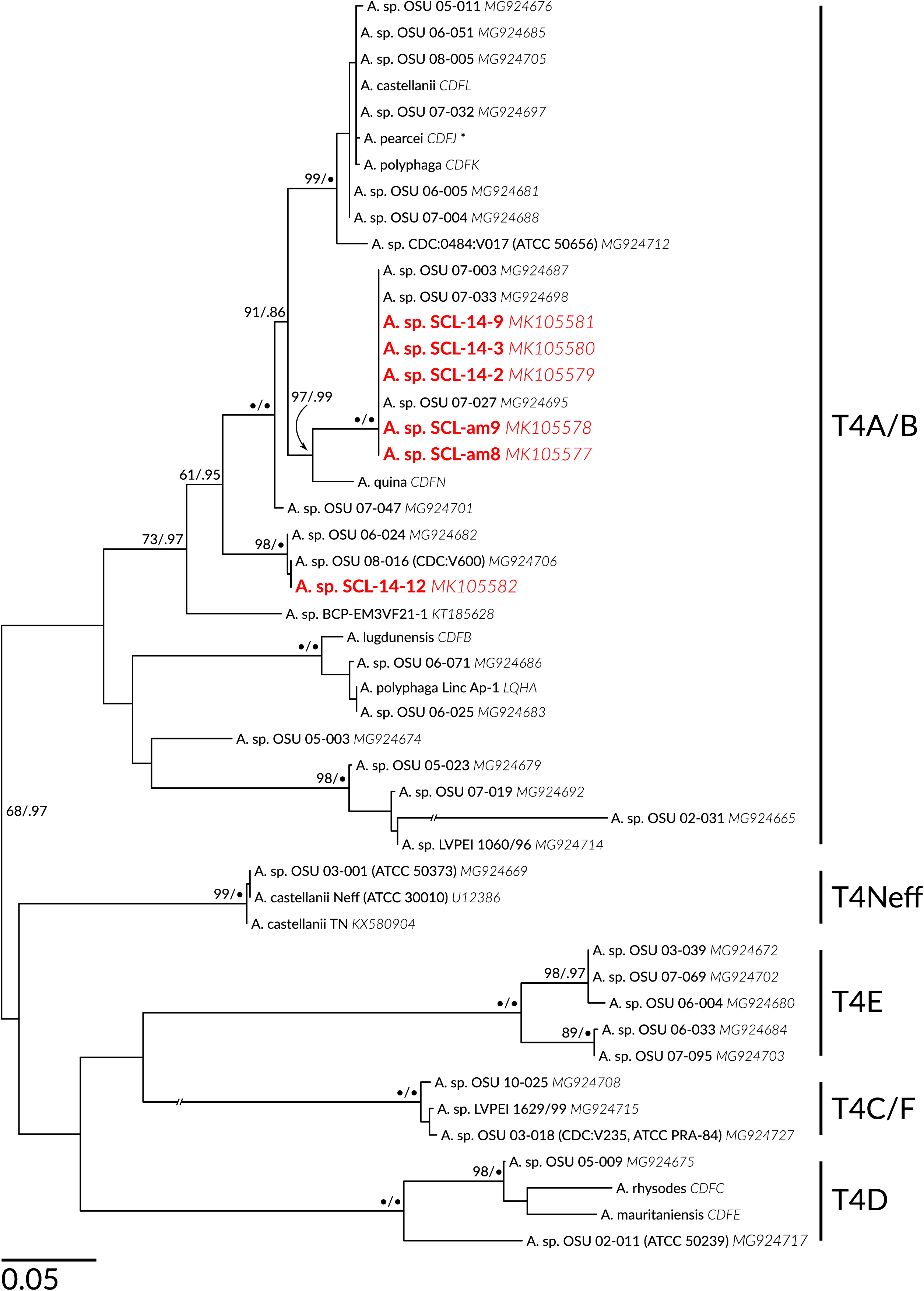
Cox1 ML-tree of *Acanthamoeba* T4 genotype rooted at midpoint. BA-tree gave the same topology. Branch supports are given as ML-bootstrap percentage / Bayesian posterior probability. • – 100% or 1.00 support. * – Identification is dubious. Support values lower than 95/.75 are not shown.

As with 16S, all permafrost strains, except SCL-14-12, possess identical sequences. Together, they form a supported clade with three sequences from the Ohio State University collection (OSU) 07-033, 07-003 (unfortunately, lacking available 18S sequences) and 07-027 (18S-T4_A_). A sequence of unknown strain identified as *Acanthamoeba quina* Pussard et Pons, 1977 (genome CDFN) appears sister to that clade with high support. Another highly supported cluster, forming a sister to the just described clade, contains the sequence of strain V017 (Rns T4_B_), sequences from genomes CDFL, CDFK, and CDFJ (probably misidentified as *Acanthamoeba pearcei* Nerad, Sawyer, Lewis et McLaughlin, 1993), and a set of OSU sequences belonging to A and B 18S subtypes. Permafrost strain SCL-14-12 groups together with strains OSU 06-024 and 08-016 (both 18S-T4_A_). One more firm cluster includes the sequence from genome CDFB, identified as *Acanthamoeba lugdunensis* Pussard et Pons, 1977. Other strains in this cluster are Linc Ap-1 (genome LQHA) and two OSU strains, both attributed to 18S-T4_A_. The last cluster in T4_A/B_ is that encompassing sequences of strain LVPEI 1060/96 (18S-T4_B_) together with OSU sequences 02-031 (18S-T4_A_), 05-023 and 07-019 (18S-T4_B_).

### Morphometrics

#### Trophozoites

The length of the measured trophozoites of all strains is between 15 and 47 μm, and the width between 5 and 27 μm (Fig. 6, Tab. 2). Locomotor forms of strain SCL-14-12 are considerably shorter (median 23 μm; □*Χ*^2^(1, 5) = 92.765, *p* < 2.2×10^-16^). The difference in length in all other strains is not significant. At the same time, the most narrow locomotor forms are those of strain SCL-am9 (median 12 μm; □*Χ*^2^(1, 5) = 51.785, *p* = 5.971×10^-10^). This leads to the significant difference in width-to-length ratio in the two strains, which is the biggest in SCL-14-12 (median 0.69) and the smallest in SCL-am9 (median 0.46) (*Χ*^2^(1, 5) = 72.762, *p* = 2.725×10^-14^). This distinction in shape could be clearly observed by eye when inspecting a series of locomotor forms (Fig. 1 A, B, C demonstrates the typical variants).

**Figure 6.**
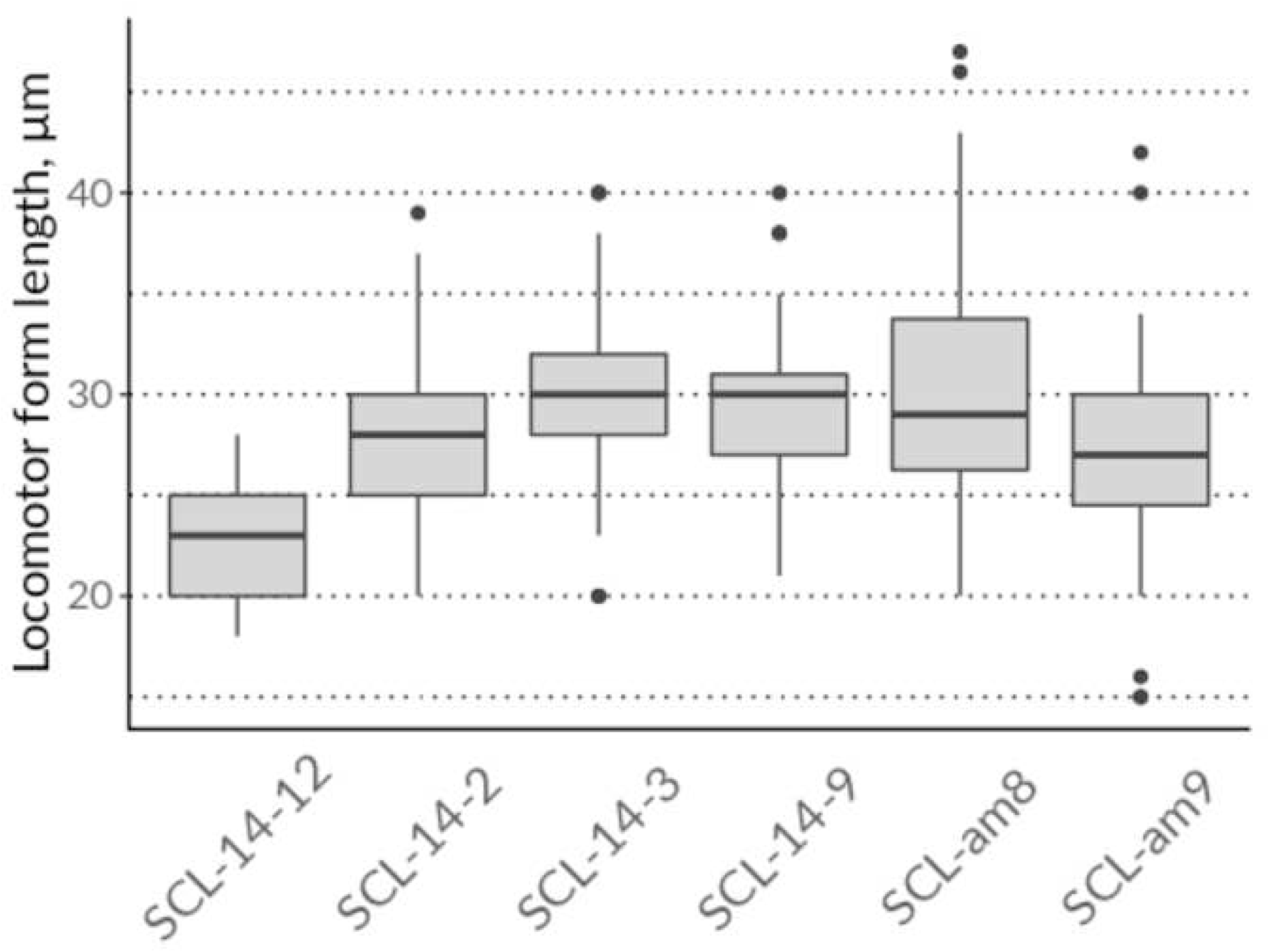
Locomotor form length in studied strains.

Some slight differences in the diameter of the nucleus and nucleolus have been observed. The median nucleus diameter in locomotor forms of strains SCL-14-12 and SCL-am9 (4 μm) are slightly lower than in other strains (5 μm) (□*Χ^2^*(1, 5) = 62.971, *p =* 2.952×10^-12^). The difference is marginally significant between SCL-14-12 and SCL-14-9, SCL-am9 and SCL-am8, and not significant between SCL-14-12 and SCL-am8. At the same time, the nucleolus in SCL-14-12 is significantly bigger (median diameter 3 μm) than in other strains (2 μm) (*Χ*^2^(1, 5) = 46.089, *p =* 8.712×10^-9^). This results in significantly higher nucleolus-to-nucleus diameter ratio in strains SCL-14-12 and SCL-am9 (not different from SCL-14-9), and also partially in strain SCL-14-9. In strain SCL-14-12, the median of the hyaline crescent breadth is the lowest (4 μm) differing significantly from all other strains (□*Χ^2^*(1, 5) = 128.58, *p* < 2.2×10^-16^). Strain SCL-14-9 has the next lowest breadth, also differing significantly from other strains at 0.05 level.

Rounded cell diameter is between 15 μm in SCL-14-12 and 19 μm in SCL-14-9 and well correlates to the cell length (Spearman’s *ρ* = 0.87, *S* = 4.6324, *p* = 0.025).

#### Cysts

We have found the cyst diameter a good character in terms of dispersion and inter-group differences. Not surprisingly, the dispersion of the cyst diameter in all estimated factor groups (“Strains” per “Culture type” design) is lower than that of the length and width in locomotor forms (MAD not greater than 12% of the median). The range of the measured cyst diameters is 10 to 24 μm (medians from 13 to 17 μm in bacterized cultures, from 14 to 17 μm in dense axenic cultures, and 15 to 19 μm in sparse axenic cultures) (Tab. 3). Permutational analysis of variance shows the significance of the effect of “Strain” (*F*(1, 5) = 263.6, *p* < .001), “Culture type” (*F*(1, 2) = 420.4, *p* < .001), and their interaction (*F*(1, 10) = 29.09, *p* < .001). Generally, the cysts formed in bacterized cultures are the smallest followed by the cysts from dense axenic cultures and then by the cysts from sparse axenic cultures, which in all strains are about the same size than the “normal” rounded trophozoites (Fig. 7), indicating almost no shrinkage during the cyst formation. In strains SCL-am8 and SCL-14-9, these differences in diameter are the most prominent, while SCL-14-12 has the smallest range (the difference is still significant).

**Figure 7.**
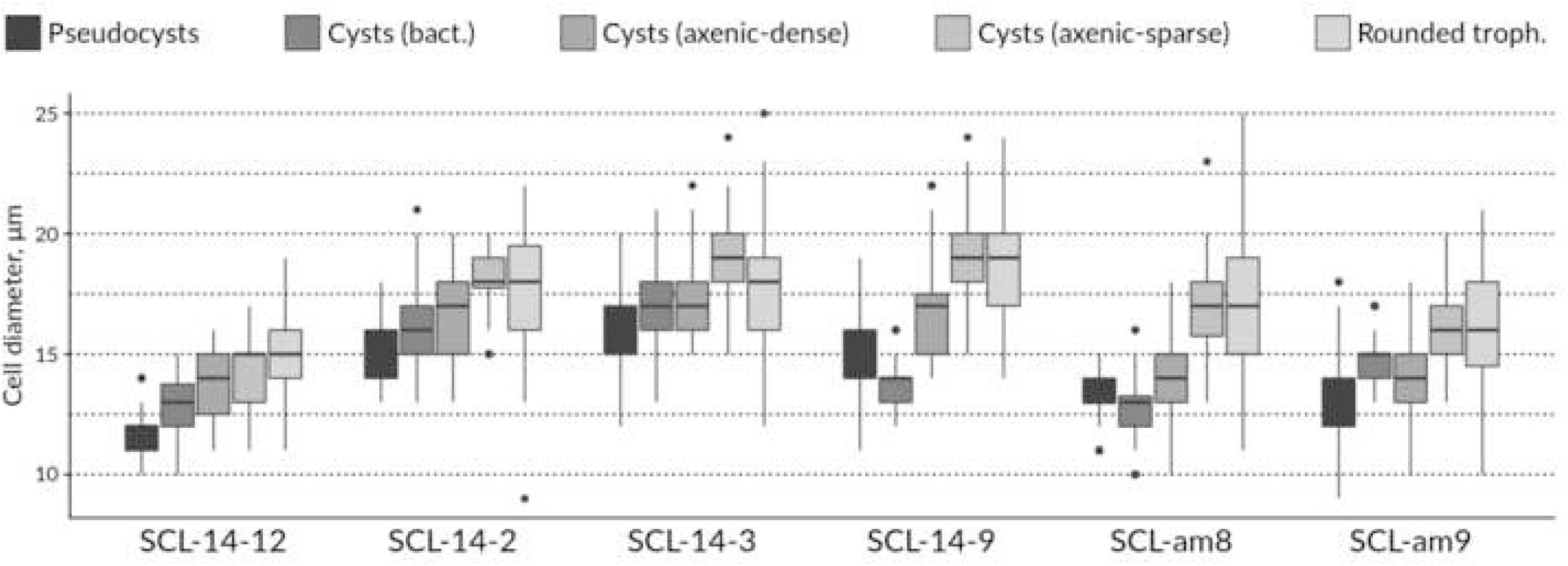
Diameter of rounded cell, cyst, and pseudocyst of the studied strains in different conditions.

No variation in the number of operculi is observed between cysts formed in different conditions.

#### Pseudocysts

The diameter of the pseudocysts in the studied strains varies from 11 to 16 μm (Tab. 4). As with cysts, SCL-14-12 is the smallest. Also, in all strains except SCL-14-9 and SCL-am8, pseudocysts are smaller than any type of cysts. In the two mentioned strains, cysts formed in bacterized cultures are slightly but significantly smaller than pseudocysts (Fig. 7).

### Growth at different temperatures

The results of the preliminary test demonstrate that none of the isolated strains can maintain the stable growth at temperatures equal to or higher than 37°C, whether as axenic cultures in PYG medium or bacterized cultures on agar plates with WG medium overlay. At 40 and 42°C, cells die entirely in about a day. At 37°C, some cells remain alive for several days (more in bacterized than in axenic cultures), but their number declined to zero to the end of the week.

32°C is not favorable for all cultures, although at this temperature, they exhibit the fastest growth (Fig. 8, Tab. 5). In strains SCL-am8, SCL-am9, SCL-14-2, and SCL-14-3, cells do not attach to the substratum and always appear roundish. A typical monolayer has never been observed. Multinucleate cells start to appear from the first counting; subsequently, their amount and size increase. The population density curve reaches stationarity very quickly. The maximum culture density at this temperature does not exceed 8×10^5^ cells mL^-1^ for SCL-am8, 5×10^5^ cells mL^-1^ for SCL-14-2 and SCL-14-2, 2×10^5^ cells mL^-1^ for SCL-am9. Strain SCL-14-12 attaches to the bottom and grows better than all in terms of the culture density (up to 2×10^6^ cells mL^-1^ have been recorded), though this constitutes a considerable decrease compared to the densities at lower temperatures. Strain SCL-14-9 demonstrates a behavior somewhat intermediate between SCL-14-12 and others. During the first two days, the culture grows well, and cells attach to the substratum. Later, most of them are found roundish and detached, and the monolayer is never formed. Multinucleate cells also appear, but later than in others and lower quantity. The density reaches 6×10^5^ cells mL^-1^.

**Figure 8.**
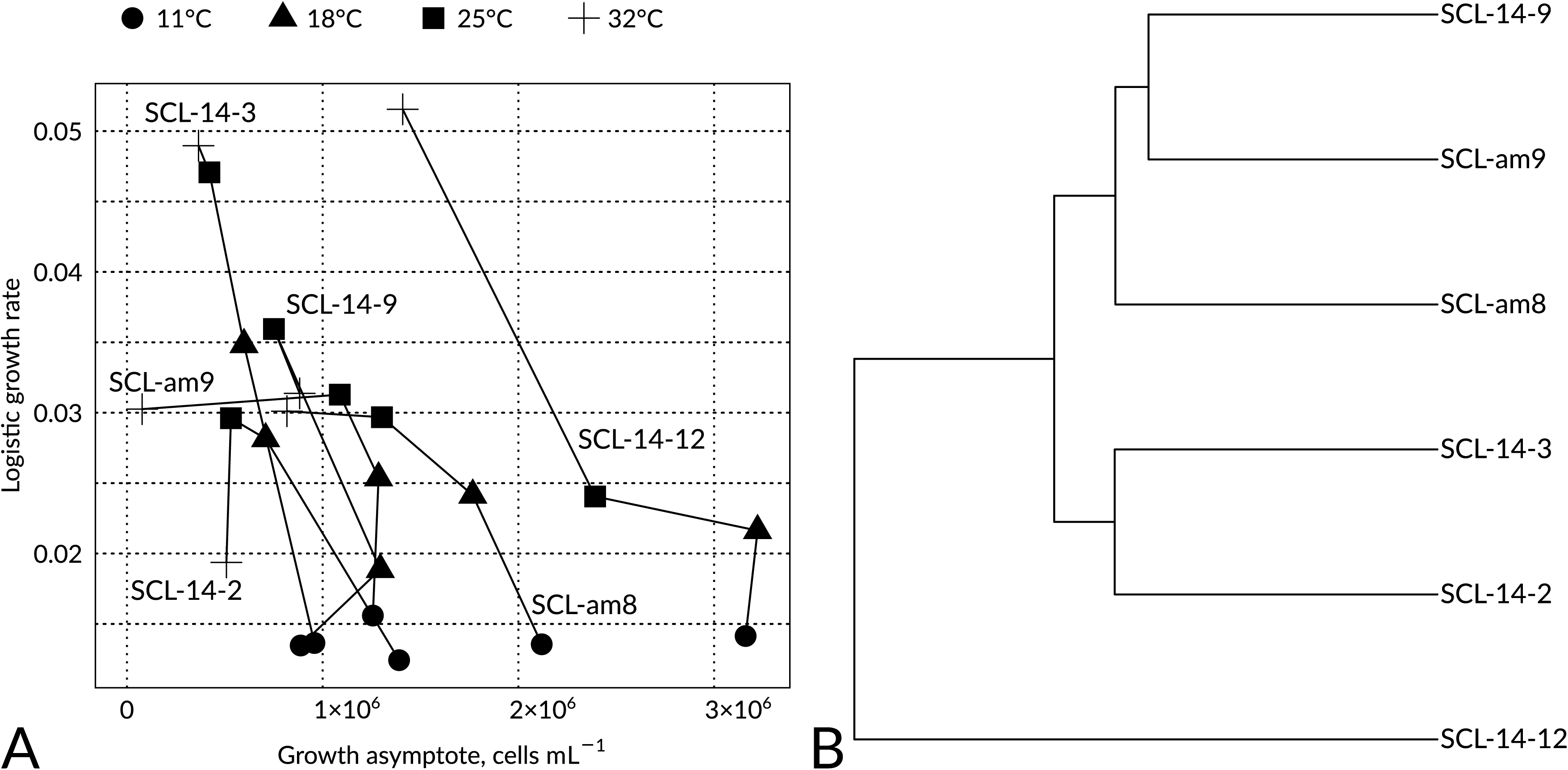
**A** – Scatter-plot representing modeled parameters of the logistic growth of the studied strains at different temperatures. Abscissa – L, ordinate – k. Lines connect points belonging to the same strain. **B** – Cluster diagram of all three logistic model parameters (scaled). Euclidean distance, Ward’s clustering method.

At 25°C, all strains grow well and form a monolayer in three to four days. Generally, the culture density increases faster than at 18°C but stops growing earlier, reaching the lower maximum. In strains SCL-am8, SCL-14-2, SCL-14-3 multinucleate cells also appear.

18°C is the optimal temperature for all strains. The maximal recorded densities are up to 8×10^5^ cells mL^-1^ in SCL-14-2 and SCL-14-3; 1.5×10^6^ cells mL^-1^ in SCL-am8, SCL-am9, and SCL-14-9; 2.8×10^6^ cells mL^-1^ in SCL-14-12) since the active growth continues longer, although the growth rate is slightly lower than at 25°C. Cultures form a monolayer in four to seven days. Cells form pseudopodia and actively move during a long time, even at the stationary phase of growth.

At 11°C, all the cultures demonstrate a considerably lower rate of growth (Fig. 4). SCL-14-12 reaches a plateau in about 600 h at a density of more than 3×10^6^ cells mL^-1^. Other strains show similar dynamics and reach a plateau in 400 h at a density of about 1.5×10^6^ cells mL^-1^ in SCL-am8 and about 1.0×10^6^ cells mL^-1^ in others.

At 4°C, all the cultures have been in a good state during the whole period of observation, judging by the cell and overall culture appearance. However, counting has shown that the cell number has been slightly declining, although dividing cells have been observed during the first two weeks.

The modeling of the logistic growth demonstrates three differences of SCL-14-12 in comparison to other strains. First, its density asymptote is higher than in other strains at any tested temperature, including 32°C. Second, its growth rate is about the same at 18 and 25°C, whereas in other strains, the growth rate at 25°C is considerably higher than at 18°C. Consequently, at 25°C, the growth rate of SCL-14-12 is lower than that of other strains. At the same time, at 32°C, the growth rate of SCL-14-12 is strikingly higher, while in others it is lower, the same, or (in SCL-14-3) just a little bit higher than at 25°C.

## Discussion

### Molecular phylogeny of *A. castellanii* complex (type T4)

#### 18S

The distinctiveness of different clades inside type T4 has begun to manifest itself since the first works on *Acanthamoeba* rDNA diversity (Booton et al., 2002; Chung et al., 1998; Fuerst et al., 2015; Gast et al., 1996; Kong, 2009; Martín-Pérez et al., 2017; Stothard et al., 1998). Subtypes T4_C_, T4_D_, T4_E_, T4_F_, and T4_Neff_, when represented by several sequences, were usually well-supported, while T4_B_ was sometimes mixed with T4_A_, and the latter mostly paraphyletic. At the same time, the relationship between the subtypes was never well-supported and appeared in part unstable. The deepest branching clade was either T4_D_, T4_E_, or a clade containing both of them. T4_Neff_ appeared sister to either T4_A/B_ or T4_D/E_ mixed clades. T4_C/F_ clade always branched after T4_D/E_, except for the phylogeny using GTR model, where T4_D_ was robustly sister to T4_A/B/Neff_ cluster (Martín-Pérez et al., 2017).

In our case, automatic alignment with subsequent filtering yields robust clades (subtypes) T4_C_ through T4_Neff_, moderately supported T4_B_, and paraphyletic T4_A_ (tree not shown). All seven subtypes as monophyletic clades could only be obtained with Jukes-Cantor substitution model, the least-likely one in MrModelTest assessment. Relationships between the subtypes are poorly supported. However, the fifth-state-coding approach demonstrates the distinctiveness of all the T4 subtypes. The fact that T4_C_—T4_F_ subtypes may be obtained from the masked alignment demonstrates the adequateness of the fifth-state-coding method used. Moreover, monophyletic and fully-supported T4_B_, T4_D_—T4_F_, and T4_Neff_ may also be obtained using the alignment of conservative regions only.

Thus, our 18S phylogeny corroborates and validates the previously obtained results. At the same time, the method bears some possible sources of bias: (i) Indels of more than one nucleotide are treated as separate events, while they may actually represent a single event, (ii) Inversions are treated as several events, (iii) The amount of transversions is probably underestimated, (iv) Unresolvable position of indels in single-nucleotide repeats. This probable bias may explain differences in topology between BA and ML trees.

#### 16S

As demonstrated by Ledee et al. (2003), 16S and 18S-based phylogenies of *Acanthamoeba* corroborate each other in terms of identified sequence types (ribotypes). This holds for T4 type as well. Further, even a restriction analysis of the 16S gene (Byers et al., 1983) has revealed remarkable heterogeneity inside 16S-T4 comparable to that found in 18S-T4. At the same time, this study has first discovered the major inconsistency between 18S and 16S phylogenies – Castellani and Ma strains (18S-T4_A_ and T4_B_, correspondingly) appeared closely related and showed differences at the level of intraspecific variation in other organisms. Other 18S-T4 type strains included in this study, namely Neff, Singh, and Page-23, were found to be as distant as other ribotypes.

The sequence-based 16S phylogeny (Ledee et al., 2003) demonstrated the identity of Castellani and Ma (as well as CDC:V042) sequences. Here, about 15% bases were removed as ambiguously aligned, which explains why differences found by Byers et al. (1983) were not observed. The rest of the sequences attributed to 18S-T4_A_ and T4_B_ subtypes were intermixed. Nine new clades were identified inside 16S-T4. Sequences of the strains attributed to 18S subtypes T4_C_, T4_D_, T4_E_, T4_Neff_ formed corresponding clades on the 16S tree (T4_F_ sequences were not included). The deepest branching clade was T4_E_ following by T4_C_, then T4_Neff_ clade. T4_D_ sequences formed a clade sister to mixed T4_A/B_ clade. Kong (2009) using a smaller dataset identified five 16S clades, namely the deepest branching T4_C_, the next branching T4_D_, then T4_Neff_, and the last branching T4_A/B_, which in turn consisted of two clusters: the one containing, among others, Ma and Castellani strains and another one containing *A. lugdunensis* L3a. Finally, Rahman et al. (2013) adding 13 T4 rns sequences obtained generally the same nine clades as Ledee et al. (2003) inside T4 subtype.

We have identified around 13 well-supported clusters inside 16S-T4 subtype. In addition to the proximity of Castellani and Ma strains, our study reveals other discrepancies between the 18S and 16S phylogenies. Permafrost strains SCL-am9 and SCL-14-9 belonging to 18S-T4_A_ subtype and SCL-am8, SCL-14-2, SCL-14-3, belonging to 18S-T4_B_ subtype, all have identical 16S sequences (even before alignment masking). Strains CDC:V042 (ATCC 50493) and CDC:V390 get into same “Rawdon” clade in 18S tree, but appear all distant in 16S tree. Strain CDC:V042 is close to Castellani strain while being placed in a different highly supported subcluster of T4_A_ by 18S phylogeny. Galka strain (ATCC 50496) groups together with SCL-am9 on the 18S tree, but is steadily apart from it on the 16S tree. By contrast, strain CDC:V390 robustly branches apart from Galka on the 18S tree but has identical 16S sequence. At the same time, strain Liu-E1 (ATCC 50709) is close to Galka in terms of 18S, and 16S sequences of these two strains are identical too. Strain SCL-14-12, which robustly clusters together with Castellani strain in absolutely all 18S analyses, steadily falls apart from it on the 16S tree. Finally, strains Rawdon (BCM:37, ATCC 50497) and CDC:V390 cluster together with high support on the 18S tree, but are steadily apart on the 16S tree.

To summarize, some of the strains demonstrate proximity of their 18S sequences and have distant 16S sequences, some, *vice versa*, have distant 18S sequences and similar 16S sequences, and other ones demonstrate similarity in both. To explain such complex dissimilarity between 18S and 16S phylogenies, three hypotheses may be involved. The first implies different rates of gene evolution in different strains, which is unlikely regarding their taxonomic proximity. As the second, the improper alignment of indels may be invoked (see the consideration of possible sources of bias above). Indeed, in single-nucleotide stretches in an alignment, indels may occur in any position and ideally should not be aligned (which is nonetheless done for the sake of parsimony). When indels are treated as the fifth state and included in the calculation, their alignment may force sequences to appear more similar than they are. This possibility cannot be completely ruled out here, but against it is the fact that only A and B and C and F 18S subtypes became intermingled in 16S tree, while the others (D, E, and Neff) stayed separate. Also, this hypothesis may explain why sequences that are close to each other in 18S phylogeny appear distant in 16S one, but not the opposite cases (as our 16S phylogeny did not account for indels).

The third hypothesis maintains that 18S subtypes T4_A_ and T4_B_, as well as T4_C_ and T4_F_, actually represent intra-genomic variants of the gene present in the cell in a different number of copies, with only the dominant one revealed by the PCR. Different strains may bear different dominant variants and thus be attributed to either A or B (C or F) subtype while sharing the same (or closely related) 16S gene sequences. Intragenomic variation in 18S gene sequence is known in different protist supergroups including other Amoebozoa (Amaral-Zettler et al., 2006; Gong et al., 2013; Gribble and Anderson, 2007; Kudryavtsev and Gladkikh, 2017; Nassonova et al., 2010; Ntakou et al., 2019; Pillet et al., 2012) and reported for several *Acanthamoeba* strains (Nuprasert et al., 2010; Qvarnstrom et al., 2013; Rahman et al., 2013; Stothard et al., 1998). Furthermore, among 18S intragenomic variants found by Rahman et al. (2013) in two strains, one belongs to T4_A_ and another to T4_B_ (the alignment is available upon request). Also, this hypothesis may explain common difficulties with obtaining a full 18S sequence from *Acanthamoeba* strains (the picture of contamination in chromatograms from a clonal culture) and rarely reported cases when different 18S variants were obtained from the same lens of a patient (Booton et al., 2002); T4_A_ and T4_C_) or from the right and left lens cases (Ledee et al., 2009); both T4_B_).

#### Cox1

The strains for which Cox1 sequences are available mostly have no full 18S sequences in the public domain. Our Cox1 phylogeny fully corroborated the 16S phylogeny in the proximity of 18S-T4_A/B_, as well as 18S-T4_C/F_, strains, the distinctiveness of other subtypes, and the internal heterogeneity of T4_A/B_ clade. Here again, strain SCL-14-12 was firmly placed into different cluster than all other permafrost strains, which suggests the correspondence of the Cox1 clusters to the 16S ones.

### Morphometrics

Traditional taxonomy of *Acanthamoeba* has been primarily based on metric (overall diameter, nucleus diameter, nucleolus diameter) and meristic (number of “angles”, or “corners”, or “arms”, i.e. protrusions of endocyst towards operculi) characters of cysts since Page’s redescription of *A. castellanii*, *A. polyphaga*, and *Acanthamoeba astronyxis* (Page, 1967). Pussard and Pons emphasized the number of angles and properties of endo- and ectocyst (smoothness, distinctiveness), which they have studied in a considerable amount of strains using different staining techniques (Pussard and Pons, 1977). However, with the number of strains growing, the variability in these characters also grew and has become overlapping. Also, as a rule, no statistical comparison of studied characters has been conducted those times, although the number of measured individuals was usually sufficient.

The use of molecular methods has shown striking similarity of some of the described species, e.g., *Acanthamoeba terricola* Pussard, 1964 and *A. castellanii* Neff strain (Visvesvara and Balamuth, 1975); *Acanthamoeba divionensis* Pussard et Pons, 1977, *Acanthamoeba paradivionensis* Pussard et Pons, 1977, *Acanthamoeba mauritaniensis* Pussard et Pons, 1977, and *Acanthamoeba rhysodes* Singh, 1952 (De Jonckheere, 1983; Liu et al., 2005). Since the beginning of the DNA sequencing era, the attention of investigators has been concentrated on molecular phylogeny, and little attempts have been made to correspond morphometric features to molecular clusters. As a result, no integrated species concept for *Acanthamoeba* that would include both molecular and morphological characters seems to exist by now.

All our strains exhibit differences in the measured morphometric characters. The morphometrics of locomotor forms demonstrates the relative stability of metric characters. For the width, the most variable trait, the median absolute deviation (MAD) is in the range from 17.4% (SCL-14-2) to 29.7% (SCL-am8) of the median (standard deviation (SD) from 16.5% to 25.5%). The MAD of the length is between 9.9% (SCL-14-3 and SCL-14-9) and 16.5% (SCL-am9) of the median (SD 12.1% to 19.4%). The level of variability found in the measured characters of cysts is even lower (not surprisingly). With the number of measured cells around 50, this level of variability allow to register significant differences (at *p* ≤ .05) even between genetically closely related strains (Tab. 2, 3).

It should be stressed that the culture conditions among the strains during the taking of measurement were held essentially the same to assure that the observed differences may be attributed to genetics. Unsurprisingly, we have found significant variability associated with the type and density of the culture in cysts (Tab. 3, Fig. 7). This indicates that the obtained results are valid for the comparison of the studied strains but may not be used to compare with literature data unless the conditions of culturing coincide exactly.

The classifications of the strains based upon different subsets of the morphometric data demonstrates different levels of similarity with molecular classification (Fig. 9). As it was discussed previously, five out of six strains are identical in mitochondrial gene sequences, so the only topological feature shared by all molecular trees is the most distant position of SCL-14-12. Not all morphological trees share this feature as well. Specifically, this is the case for locomotor form and sparse axenic cyst measurements (Fig. 9F, G), which also drives the overall configuration (Fig. 9D). The dense axenic cyst and bacterized cyst data (Fig. 9H, I), as well as the pseudocyst data, do not show this topology. Notably, monoclonal bacterized culture on an agar plate is the most common setting in which measurements have been taken from different *Acanthamoeba* strains, partially because not in every case a culture has been axenized, and partially because this has been (probably rightly) believed to most closely resemble the natural conditions. Our results suggest that measurements taken in axenic conditions may better reflect genetic differences.

**Figure 9.**
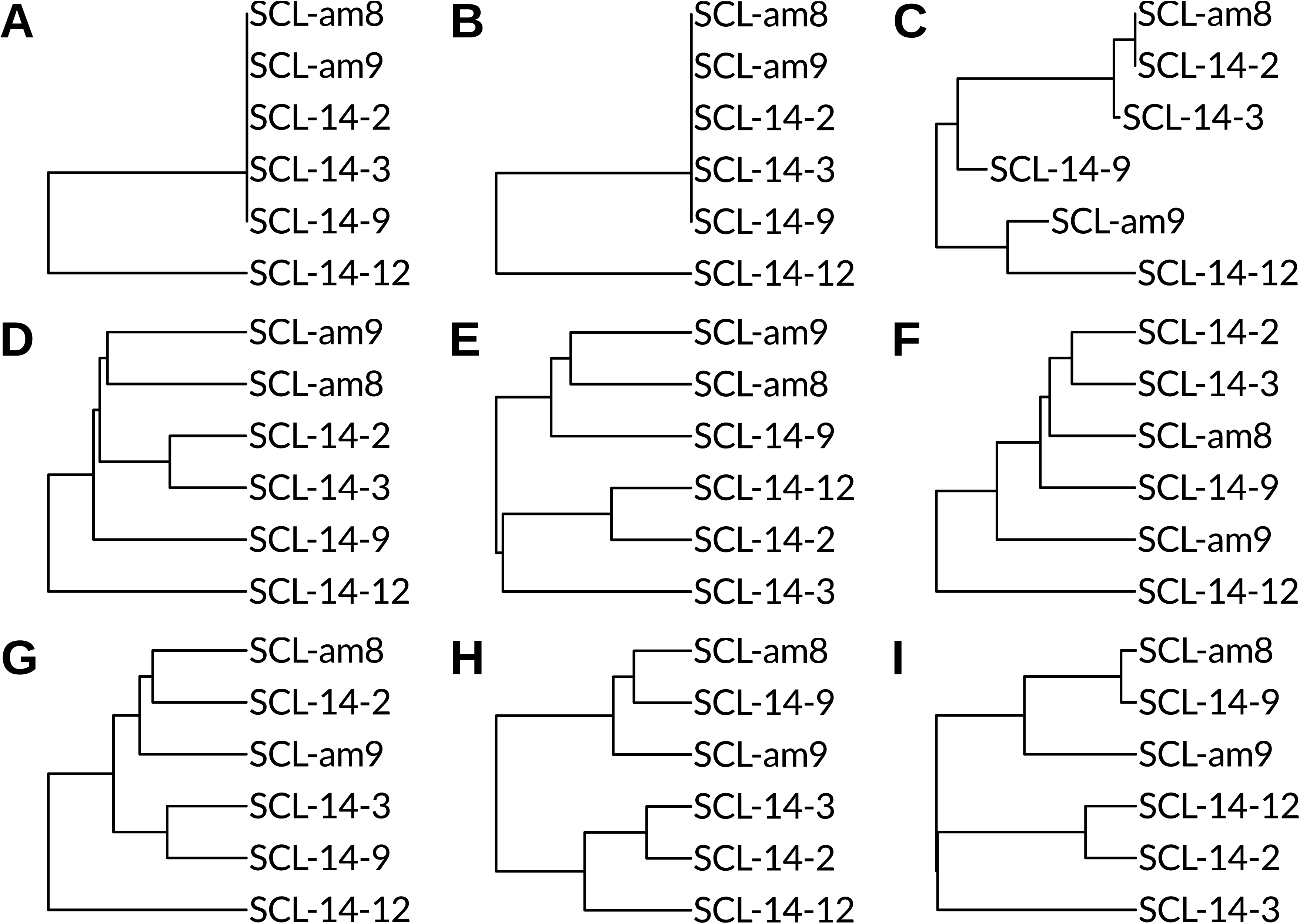
Classification of the strains based on different data. **Gene trees: A** – Cox1 NJ tree, observed distances; **B** – 16S NJ tree, observed distances; **C** – 18S tree extracted from the complete tree of T4 type. **Morphometric trees: D** – All features included; **E** – Cysts and pseudocysts only; **F** – Trophozoites only; **G** – “Sparse axenic” cysts; **H** – “Dense axenic” cysts; **I** – “Bacterized culture” cysts. All trees are rooted at midpoint. Not to scale.

Thus, the most distant position of strain SCL-14-12 is not absolutely stable when different sets of features and different culture conditions are compared. At the same time, we tend to attach greater importance to the measurements of locomotor forms as to embracing more different cell features. In addition to significant differences in linear dimensions, strain SCL-14-12 also differs from others by its proportions, specifically the width to length, nucleolus to nucleus diameter, and hyaloplasm width to cell length ratios. It should be noted that not only SCL-14-12 is the most distant strain in the locomotor form-based classification, but also the further branching (SCL-am9, then SCL-14-9) partially follows the order of the 18S tree (Fig. 9C).

So, to reliably compare morphometric data between strains in *Acanthamoeba* (and other amoebae), the establishment of standardized culture conditions is absolutely necessary. Axenic culture in a defined medium is the best candidate for such a standard, but the temperature should obviously be considered as well.

### Growth at different temperatures

As with the morphometric data, we observe differences in the culture growth are between all studied strains and temperatures. Generally, the higher the temperature, the higher the logistic growth rate, and the lower the culture density asymptote. Also, at temperatures from 25°C and higher, multinucleate cells appear, prevailing over the normal mononucleate cells at 32°C in most strains. James and Byers (1967) also observed multinucleates in aging cultures; their amount was increased by agitation or air bubbling. Corsaro et al. (2015) observed multinucleates in a culture of *Acanthamoeba micheli* Corsaro, Walochnik, Köhsler et Rott, 2015. Most likely, multinucleate cells result from incomplete cell division (karyokinesis not followed by cytokinesis), but the possibility of cell fusion also cannot be ruled out (Pigon, 1972). Our data strongly supports the first way.

In the growth data, strain SCL-14-12 is also obviously different from others. At 25°C, it shows approximately the same growth rate as at 18°C, while other strains increase it. At 32°C, no multinucleate cells appear in this strain, and the cells are attached to the substratum. Also, the growth rate is the highest. Thus, in addition to higher growth limits at all temperatures tested (a feature stemming probably from its smaller size), SCL-14-12 is characterized by a broader range of temperatures suitable for normal culture growth.

The potential pathogenicity of *Acanthamoeba* strains is often addressed by testing their possibility to grow at 34°C, a temperature of the human cornea (Walochnik et al., 2000), and 37°C. All permafrost strains except SCL-14-12 demonstrate clear signs of dysfunction already at 32°C, and may thus be considered non-pathogenic. For SCL-14-12, the potential to cause keratitis cannot be excluded.

## Conclusion

Although a complex comparative study of strains inside *A. castellanii* complex (ribotype T4) remains a future perspective, our investigation demonstrates that the results of consistently applied morphometrics and ecophysiological studies (characteristics of growth at different temperatures) are in line with molecular variability of several kinds of genes (nuclear and mitochondrial) and may thus form a basis for *Acanthamoeba*-specific species concept. Amid the diversity of the entire genus, all studied strains are closely related to each other. However, strain SCL-14-12 is unique in all three sequenced genes, while the others possess the same sequences of the mitochondrial genes. Phylogenies built with all available sequences firmly place SCL-14-12 apart from the other strains. At the same time, it differs by many morphometric characters of both trophozoites and cysts, including proportions, as well as growth characteristics at different temperatures. Besides, directly observable features of cultures also distinguish this strain from the others. Other strains also exhibit some variation (and are obviously different in their 18S sequences), but this variation does not involve all studied features and is smaller in magnitude. Taking all this into account, we consider strain SCL-14-12 and other permafrost strains as two separate species. This conclusion may also be extrapolated onto other clusters obtained in mitochondrial phylogenies inside T4_A/B_ clade, if only morphological and ecophysiological differences of strains between those clusters are consistent, which needs further investigation. Until such investigation placing phenotypic divergence in the framework of steady multigene phylogeny is done, a revision of *A. castellanii* species complex and thus the genus as a whole seems not possible.

Although most of *Acanthamoeba* 18S rDNA types are conceivably real taxa, in *A. castellanii* complex, the phylogenetic signal of this gene is biased, and thus it should not be used for species delimitation in this group. Mitochondrial gene phylogenies seem to be more suitable for this purpose.

## Acknowledgements

This study partially utilized microscopy equipment of the Core Facility Centres “Development of molecular and cell technologies” and “Culture Collection of Microorganisms” of Saint Petersburg State University (SPSU). We thank Dr. Alexei Smirnov for providing the possibility to use these facilities. We are obliged to all Soil Cryology lab members who participated in field sampling. We would also like to thank Prof. Dr. Jean-Michel Claverie, Laboratoire Information Génomique et Structurale, Aix Marseille Université, for his valuable comments on the manuscript.

This study was conducted in the framework of the Governmental Program 0191-2019-0044 and supported by Russian Foundation for Basic Research (RFBR) grants 18-04-00824 (isolation and culture establishment) and 17-54-150003 (description of the strains).

