## Supplementary for "Isolates from ancient permafrost help to elucidate species boundaries in *Acanthamoeba castellanii* complex (Amoebozoa: Discosea)"

### Supplementary material

to the journal article 'Isolates from ancient permafrost help to elucidate species boundaries in Acanthamoeba castellanii complex (Amoebozoa: Discosea)' by S. Malavin and L. Shmakova

#### Material and Methods

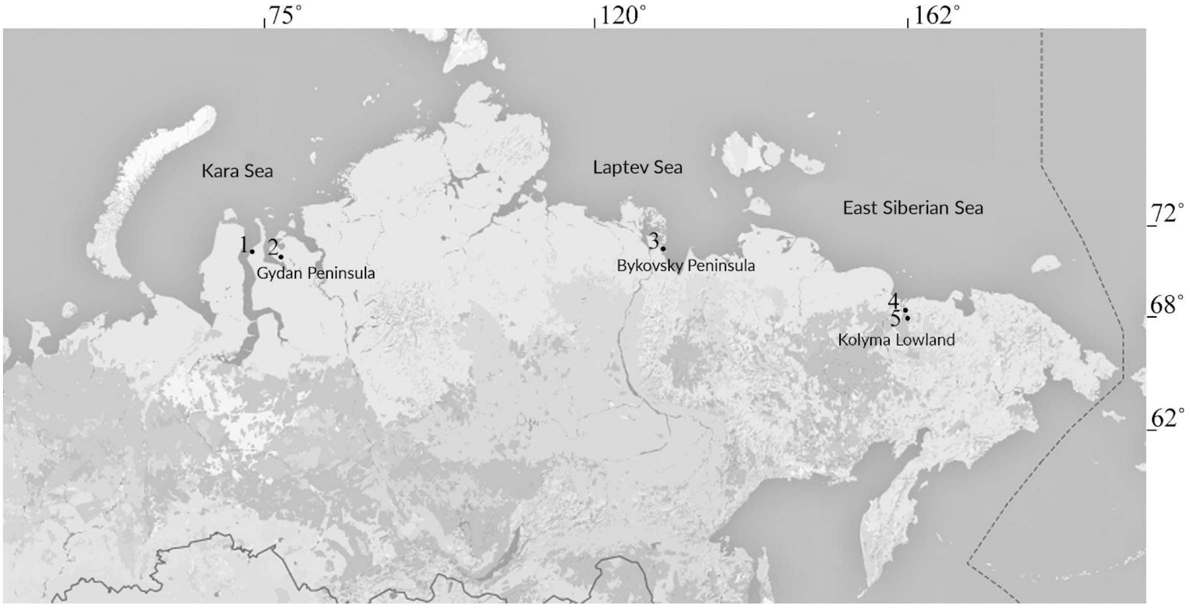


**Supplementary Figure 1** Sampling sites for the obtained Acanthamoeba strains.
**1** SCL-14-3; **2** SCL-14-2, SCL-14-9; **3** SCL-am9; **4** SCL-am8; **5** SCL-14-12.

#### Detailed description of sequence acquisition

The almost complete nuclear SSU rRNA gene (18S) was amplified using eukaryotic primers RibA and RibB (Supplementary Tab 1). The 25 µL PCR mixture contained 0.5 µL (0.2 µM) of each primer, 0.5 µL (0.2 µM) of each dNTP, 0.75 µL (1.5×) of Encyclo Polymerase (blend of hot-start Taq and high-fidelity polymerases; Evrogen, Russia), and 2.5 µL of Encyclo buffer (3.5 µM $MgCl_2$). The amount of DNA template measured in 2 µL by Qubit fluorometer (Invitrogen, USA) varied from 30 to 80 ng per reaction. After the initial denaturation at 96°C for 3 min, the reaction was cycled 35 times and finished with 1 min final elongation at 55°C and 9 min at 72°C. Each cycle consisted of 20 sec at 96°C, 30 sec at 55°C, and 2 min 20 sec at 72°C.

The mitochondrial SSU rRNA gene (16S) was amplified with primers mt1 and mt1541 (Supplementary Tab 1) flanking an almost complete gene (Ledee et al., 2003). The reaction volume, mixture component concentrations, and cycling conditions were the same as for the 18S gene, except for 2-min elongation.

The 5’-fragment of mitochondrially encoded cytochrome c oxidase subunit I gene (Cox1) 674 to 741 bp in length was amplified using universal primers LCO1490 and HCO2198 (Folmer et al., 1994). The reaction volume, mixture component concentrations, and cycling conditions were as for RNA genes, except for 40-sec elongation.

All amplified fragments were gel-purified and Sanger-sequenced on an automated DNA sequencer in Evrogen Co. (Russia). Additional primers used for sequencing are listed in Supplementary Table 1, and their approximate positions schemed in Supplementary Figure 1.

**Supplementary Table 1**. Primers used in the study

| Name | Sequence 5'-3' | Direction | Gene | Purpose | Reference |
|---|---|---|---|---|---|
| RibA | ACCTGGTTGATCCTDCCAGT | fw | 18S | Ampl., seq. | Medlin et al., 1988 |
| RibB | TGATCCTTCTGCAGGTTCAC CTAC | rev | 18S | Ampl., seq. | Medlin et al., 1988 |
| P1fw | CAAGTCTGGTGCCAGCAGC | fw | 18S | Seq. | Walochnik et al., 2004 |
| P1rev | GCTGCTGGCACCAGACTTG | rev | 18S | Seq. | Walochnik et al., 2004 |
| s12.2 | GATCAGATACCGTCGTAGTC | fw | 18S | Seq. | |
| s12.2R* | GACTACGACGGTATCTRATC | rev | 18S | Seq. | |
| P3fw | CAGGTCTGTGATGCCCTTAG | fw | 18S | Seq. | Walochnik et al., 2004 |
| P3rev | CTAAGGGCATCACAGACCTG | rev | 18S | Seq. | Walochnik et al., 2004 |
| mt1 | TTGTATAAACAATCGTTGGG T | fw | 16S | Ampl., seq. | Ledee et al., 2003 |
| mt1541 | AAAATTTTGTCCAGCAGCA | rev | 16S | Ampl., seq. | Ledee et al., 2003 |
| mt600 | AAGTGTAAAGGTGAAATT | fw | 16S | Seq. | Ledee et al., 2003 |

* The degenerate primer s12.2R was used not with any particular purpose, but simply due to availability.

**Supplementary Table 1** (continued)

| Name | Sequence 5'-3' | Direction | Gene | Purpose | Reference |
|---|---|---|---|---|---|
| mt900 | CAAATTAAACCACATACT | rev | 16S | Seq. | Ledee et al., 2003 |
| LCO1490 | GGTCAACAAATCATAAAGA TATTGG | fw | CO1 | Ampl., seq. | Folmer et al., 1994 |
| HCO2198 | TAAACTTCAGGGTGACCAAA AAATCA | rev | CO1 | Ampl., seq. | Folmer et al., 1994 |

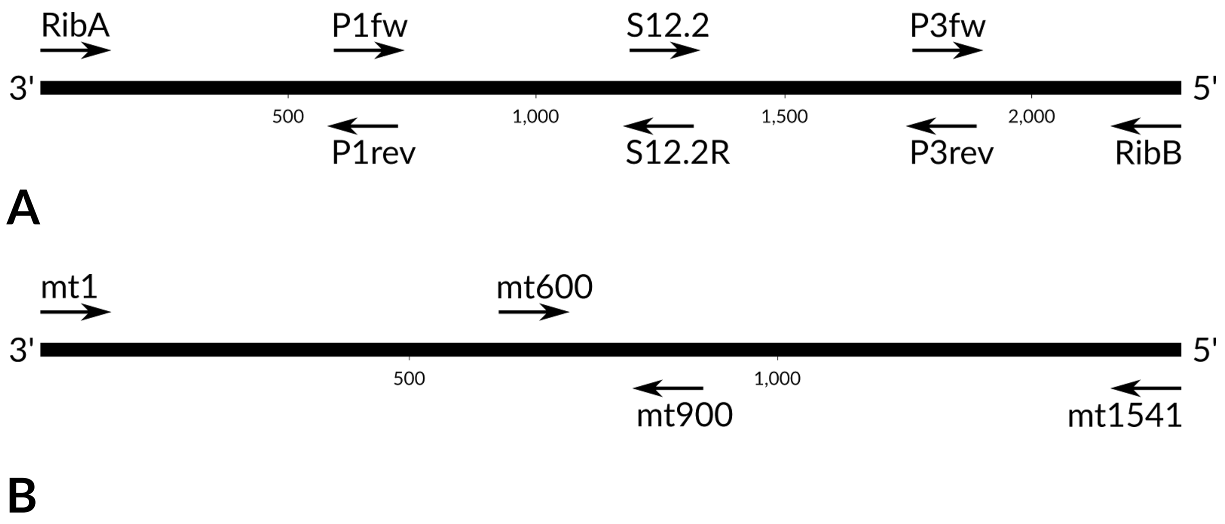


**Supplementary Figure 2** Primers used in this study. **A** Amplification and sequencing of 18S SSU rDNA; **B** Amplification and sequencing of 16S SSU rDNA (mitochondrial). Shown nucleotide coordinates are approximate

#### Detailed description of phylogenetic analysis

In all the analyses described below, sequences were aligned by MAFFT v. 7.307 L-INS-i algorithm (Katoh and Standley, 2013) with default scoring matrix and gap penalties; the maximum iteration number was set to 1000. The alignment was further polished manually. Introns present in some sequences were deleted. The phylogeny was reconstructed by Bayesian analysis of posterior probabilities (Bayesian inference, BI) and maximum likelihood (ML) approaches. Substitution models were selected by MrModeltest v. 2.4 (Nylander, 2018) using hierarchical likelihood ratio test (hLRT) and standard Akaike information criterion (AIC). If not indicated, both BI and ML trees had the same topology. BI was performed in a parallelized version of MrBayes v. 3.2.6 (Ronquist et al., 2012) with 3 parallel independent runs of 1 cold and 4 heated chains. The temperature parameter was 0.1

unless indicated. Tree populations were sampled from each 500th generation. The total number of generations is indicated separately for each analysis. The convergence of runs was assessed using the built-in MrBayes tools; a proportion of generations discarded as burn-in is reported below in each case. ML analysis was done in IQTree v. 1.6.6 (Nguyen et al., 2015). A consensus tree was built using ultrafast bootstrap approximation with 1000 replicates and a hill-climbing nearest-neighbor interchange (NNI) optimization of each bootstrap tree (Hoang et al., 2018). All alignments are available from the authors upon request.

18S *Acanthamoeba* sequences with length equal to or greater than 2 kb were retrieved from GenBank with the following search query: 'Acanthamoeba[ORGN] AND (18S OR SSU OR "Small subunit ribosomal") AND 2000:3300[SLEN]'. Additional *Acanthamoeba* sequences deposited under different names were revealed by blasting several sequences from the obtained collection.

Several sequences were then removed from the alignment due to either the presence of long obviously artifactual gaps or high similarity to others. Small gaps (not greater than 5 nt) in highly conserved regions (the same motif in all sequences) were considered artifactual and manually filled with the same motif to increase the number of estimated sites. The resulting alignment was filtered by BMGE 1.12 with the default region size and similarity matrix. BI and ML analysis were done with GTR+Γ4+I substitution model (general time-reversible model with the discrete 4-category gamma approximation of the rate variation across sites and a correction for invariable sites). In BI, $5\times10^6$ generations of trees were produced. Temperature parameter for the heated chains was set to 0.05 as the default 0.1 did not provide optimal swapping. In ML and BI analyses, previously identified genotypes were generally highly supported, although their relationship remained unresolved and differed depending on the stringency of alignment trimming (trees not shown). All permafrost isolates robustly grouped within T4 genotype, which always appeared monophyletic. We thus concentrated on its internal structure.

To benefit from the structure of gappy hypervariable regions, gaps were re-coded as following. T4 sequences were selected from the overall alignment (before filtering), and alignment was divided into two partitions: conservative regions (well-aligned automatically) and variable regions (not aligned; see fig. 5). In the variable-region partition, first, all purines and pyrimidines were aligned. Transversions were only allowed if at least two nucleotides surrounding the questioned site aligned well across all or most of the sequences. Next, individual nucleotides were aligned inside the strips of purines and pyrimidines so as to minimize the number of different patterns inside the strip. If obvious two- or several-nucleotide inversion events were observed, those were allowed. Insertions observed in one or a few sequences only (mostly nucleotide duplications) were removed to reduce the number of sites, and also because they may represent sequencing bias. Single-nucleotide stretches of different width were reduced, so as none of the sequences differed from each other by more than 1 nucleotide, again to reduce the number of sites and shorten the branches at the same time. After that, constant sites were removed, and the resulting 154 variable sites were re-coded as morphological characters with gaps represented as a fifth character state.

The analysis of the partitioned dataset was conducted with GTR+Γ4+I substitution model for the conservative-sites partition (BI: $2 \times 10^6$ generations, temperature 0.03) and JC+Γ4+I model for the variable-sites partition ("MK+G4+I+ASC" in IQTree, where ASC is an ascertainment bias correction model (Lewis, 2001). The partitions were allowed to have separate proportions of invariable sites and branch length sets.

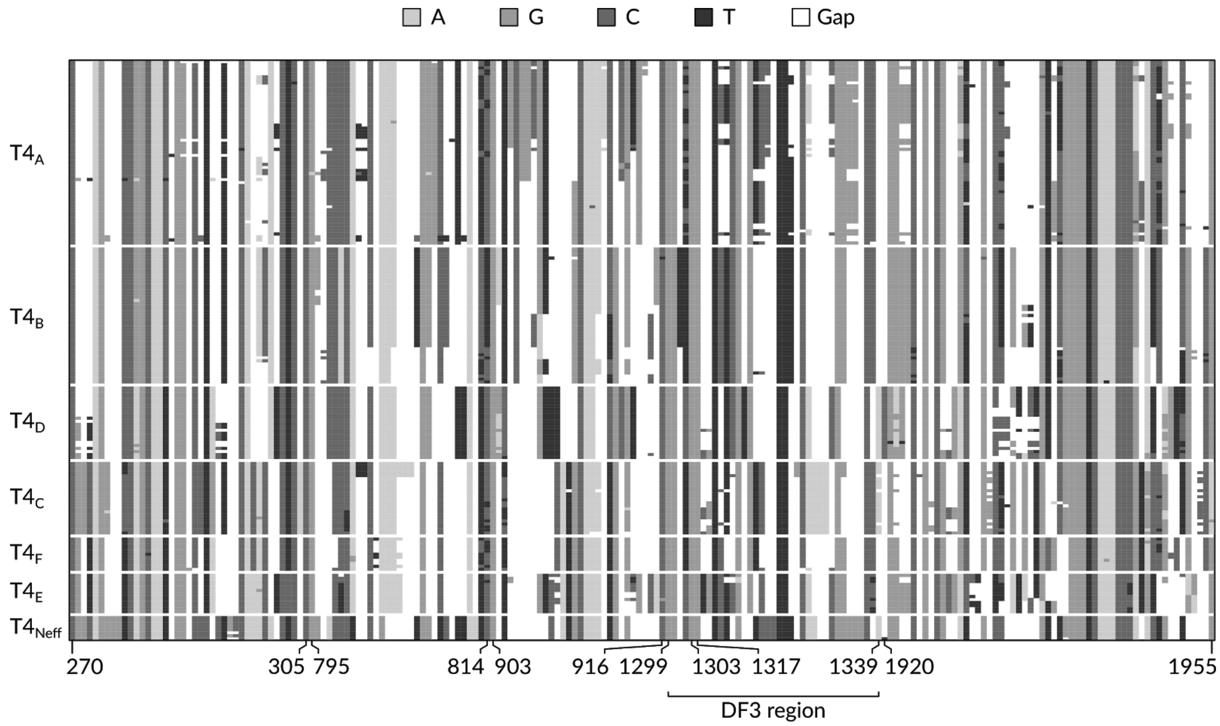


**Supplementary Figure 3** Alignment of concatenated hypervariable regions (partition 2) of the studied "full-length" T4 18S sequences. Numbers indicate locations in *Acanthamoeba sp.* Neff GenBank acc. U07416.

16S sequences were retrieved from GenBank by a search query 'Acanthamoeba[ORGN] AND 16S[TITLE] AND 1000:2000[SLEN]'. The alignment was trimmed by BMGE with the default parameters. Corresponding full or partial 18S sequences were searched, and in case of their availability, the 18S-genotype was attributed (for partial sequences, the attribution was primarily made by the structure of the hypervariable region in DF3 gene fragment). The phylogeny was reconstructed with GTR+I+Γ4 model (BI: $2 \times 10^6$ tree generations).

Cox1 sequences were obtained from GenBank by searching for 'Acanthamoeba[ORGN] AND (COI OR Cox1 OR "cytochrome c oxidase")'. 18S genotyping was done as for 16S sequences. Only T4-genotype sequences were selected. Additional Cox1 sequences were retrieved by blasting SCL-am8 sequence against draft *Acanthamoeba* genomes available in GenBank with the default parameters. The phylogeny was reconstructed with HKY+I+Γ4 model (HKY+I+F+Γ4 in IQTree) (BI: $10^6$ generations).
